# Defining the roles of pyruvate oxidation, TCA cycle, and mannitol metabolism in methicillin resistance *Staphylococcus aureus* catheter-associated urinary tract infection

**DOI:** 10.1101/2022.10.28.514332

**Authors:** Santosh Paudel, Sarah Guedry, Chloe LP Obernuefemann, Scott Hultgren, Jennifer N Walker, Ritwij Kulkarni

**Affiliations:** Department of Biology, University of Louisiana at Lafayette, Lafayette, LA-70504; Center for Women’s Infectious Disease Research, Department of Molecular Microbiology, Washington University School of Medicine, St. Louis, MO-63110; Department of Microbiology and Molecular Medicine, McGovern Medical School, University of Texas Health Science Center at Houston, TX-77030; Department of Epidemiology, Human Genetics, and Environmental Science, School of Public Health, University of Texas Health Science Center at Houston

**Author notes:** Corresponding author **email:**, **Address:** 410 E. St. Mary Blvd, Billeaud Hall Room 108, Department of Biology, University of Louisiana at Lafayette, Lafayette, LA-70504.

## Abstract

Methicillin resistant *Staphylococcus aureus* (MRSA) is an important cause of complicated urinary tract infection (UTI) associated with the use of indwelling urinary catheters. Previous reports have revealed host and pathogen effectors critical for MRSA uropathogenesis. Here, we sought to determine the significance of specific metabolic pathways during MRSA UTI. First, we identified 16 mutants from the Nebraska transposon mutant library in the MRSA JE2 background with significantly reduced growth in pooled human urine (HU). Among these, five genes targeted by transposon mutation also showed significant upregulation upon exposure to HU for 2 h. This prompted us to generate transposon insertion mutants in the uropathogenic MRSA 1369 strain that were defective in TCA cycle (Δ*sucD*, Δ*fumC*), mannitol metabolism (Δ*mtlD*), and pyruvate oxidation and branched chain fatty acid synthesis (Δ*lpdA*). Compared to the WT, the Δ*lpdA* mutant showed a significant defect growth in HU and colonization of the urinary tract and dissemination to spleen in the mouse model of catheter-associated UTI (CAUTI), which may be attributed to its increased membrane hydrophobicity and higher susceptibility to killing in blood. MRSA 1369 Δ*sucD*, Δ*fumC*, and Δ*mtlD* mutants were not defective for *in vitro* growth in HU but showed significant fitness defects in the CAUTI mouse model. Overall, identification of novel metabolic pathways important for the urinary fitness and survival of MRSA can be used for the development of novel therapeutics.

**Importance:** While *Staphylococcus aureus* has historically not been considered a uropathogen, *S. aureus* urinary tract infection (UTI) is clinically significant in certain patient populations, including those with chronic indwelling urinary catheters. Moreover, most *S. aureus* strains causing catheter-associated UTI (CAUTI) are methicillin-resistant *S. aureus* (MRSA), which is difficult to treat as it limits treatment options and has the potential to deteriorate into life-threatening bacteremia, urosepsis, and shock. In this study, we found that pathways involved in pyruvate oxidation, TCA cycle, and mannitol metabolism are important for MRSA fitness and survival in the urinary tract. Improved understanding of the metabolic needs of MRSA in the urinary tract may help us develop novel inhibitors of MRSA metabolism that can be used to treat MRSA-CAUTI more effectively.

## INTRODUCTION

Historically, *Staphylococcus aureus* was considered an atypical uropathogen; however recent reports highlight its clinical significance in the urinary tract (UT) (1-3). These reports indicate *S. aureus* is an increasing cause of asymptomatic bacteriuria and complicated urinary tract infections (UTIs) primarily in the elderly, those with recent hospitalization, and in individuals with indwelling urinary catheters (1, 2, 4-7). Moreover, *S. aureus* UTI has become a clinical concern over the last two decades due to the increasing incidence methicillin-resistant *S. aureus* (MRSA) detected in human urine (HU), indicating that the urinary tract can be a reservoir for drug resistance (6, 8, 9). Additionally, *S. aureus* UT colonization is a known precursor for life-threatening invasive infections such as bacteremia, urosepsis, and shock (6, 9-11). The role of urinary catheterization in predisposing individuals to MRSA UT colonization has also been validated in the C57BL/6 mouse model of catheter-associated UTI (CAUTI) (12). In our previous study, catheterized mice showed 300-fold higher MRSA bladder burdens at 1 day post infection (dpi) compared to their non-catheterized counterparts and MRSA persistence was marked by significantly higher bacterial burden in the bladder and kidneys of catheterized mice at 14 dpi (12).

The UT is a moderately oxygenated microenvironment containing urine, which is a high osmolarity, iron-limiting, dilute mixture of amino acids and peptides (13). The successful colonization of the UT is contingent on the ability of uropathogens to transport amino acids and peptides across the membrane and catabolize them via the TCA cycle and gluconeogenesis. Indeed, mutants of uropathogenic *Escherichia coli* (UPEC) ablated for peptide transport as well as mutants of UPEC and *P. mirabilis* deficient for TCA cycle and gluconeogenesis exhibited significantly reduced fitness in the mouse model of ascending UTI (14, 15). In our previous study, 2 h-long exposure of uropathogenic MRSA 1369 to HU induced the expression of oligopeptide transporters (oppBCDFAA) and enzymes catalyzing the TCA cycle and gluconeogenesis (16). In the current study, we provide evidence to further support a central role for amino acid catabolism in MRSA uropathogenesis. Our screen of the *S. aureus* Nebraska transposon mutant library (JE2 strain background) identified 16 mutants that showed normal (WT-like) growth in nutrient rich BHI but displayed a growth defect in HU. Next, we generated select metabolic gene mutations in the uropathogenic strain MRSA 1369, confirmed their growth defects in HU, and competed these mutants with the WT to test *in vitro* growth in HU and *in vivo* colonization of the mouse model of CAUTI. Compared to the MRSA 1369 WT, TCA cycle enzyme mutants Δ*fumC* (fumarase) and Δ*sucD* (succinyl coA synthase) and Δ*mtlD* (mannitol-1-phosphate dehydrogenase) did not display growth defects in HU, however, all three showed modest competitive disadvantages in a mouse model of CAUTI. Notably, MRSA 1369Δ*lpdA* – deficient for the activity of dihydrolipoamide dehydrogenase, a pyruvate dehydrogenase complex enzyme – was significantly defective for growth and fitness in HU, as well as for the colonization of bladder and kidney tissues in a mouse model of CAUTI. Furthermore, the poor *in vivo* survival of MRSA 1369Δ*lpdA* may be attributed to increased hydrophobicity and susceptibility to killing in blood. This work supports previous reports that LpdA activity regulates the biosynthesis of branched chain fatty acids resulting in reduced membrane fluidity and increased cell surface hydrophobicity (17), that *S. aureus* Δ*lpdA* mutants do not produce surface protein A, a critical virulence factor (18), and that *S. aureus* Δ*lpdA* mutants exhibit poor survival in a mouse model of systemic infection (19).

In summary, these results improve our knowledge of how MRSA may utilize specific metabolic pathways for survival and proliferation in the host UT. Our observations may also be applied towards the development of novel therapeutics against MRSA UTI, a clinically relevant endeavor in the face of an ever-shrinking repertoire of effective antibiotics and against an expanding pool of emergent drug-resistant strains of *S. aureus*.

## Materials and Methods

### Bacterial strains, HU, and reagents

The WT MRSA strain JE2 and the Nebraska transposon mutant library (NTML), which consists of 1920 mutant strains generated by the insertion of mariner transposon *bursa aurealis* in the genome of MRSA JE2, were obtained from the network on antimicrobial resistance in *Staphylococcus aureus*, NARSA (20, 21). The NTML strains and WT JE2 were screened for growth in HU alongside the uropathogenic *S. aureus* strain MRSA 1369 (12, 16). The genes targeted by select transposon insertion mutants were moved into the MRSA 1369 background via phage 11 transduction following previously published methods (12) and are listed in **Table 1**. For simplification, these transposon mutants are referred to as Δ*gene_name*. The WT and mutant strains were cultured overnight at 37°C and shaking at 200 rpm in brain heart infusion (BHI) broth. Overnight cultures of bacterial strains were centrifuged, washed once in sterile Dulbecco’s phosphate-buffered saline (D-PBS) at room temperature, resuspended in D-PBS, and used for in vitro experiments.

**Table 1.** Strains used in this study.

| Strain | Description | Source |
| --- | --- | --- |
| JE2 | Wildtype background strain in which the NTML was generated. | (20, 21) |
| NTML | Collection of 1920 transposon mutants in the JE2 background. | (20, 21) |
| JE2 *sucD*::tn | Transposon mutant in *sucD* | This study |
| JE2 *fumC*::tn | Transposon mutant in *fumC* | This study |
| JE2 *mltD*::tn | Transposon mutant in *mltD* | This study |
| JE2 *lpdA*::tn | Transposon mutant in *lpdA* | This study |
| MRSA 1369 | Urinary tract clinical isolate | (12) |
| MRSA 1369 Δ*sucD* | Transposon mutant in *sucD* | This study |
| MRSA 1369 Δ*fumC* | Transposon mutant in *fumC* | This study |
| MRSA 1369 Δ*mltD* | Transposon mutant in *mltD* | This study |
| MRSA 1369 Δ*lpdA* | Transposon mutant in *lpdA* | This study |

HU collected from healthy female volunteers between age 18 to 45 years (protocol IRB-22-054-BIOL approved by IRB, UL Lafayette) was filter sterilized through a 0.22-μm filter, aliquoted, and stored at -20°C. At the time of experiment, urine aliquots from at least 3 separate donors were pooled.

### Screening of NTML strains and WT for growth in HU

The 1920 MRSA JE2 mutants were screened in vitro for growth in HU alongside WT JE2 and MRSA 1369. Similar inocula of a mutant or WT were mixed with pooled HU and 200 μl was added to a 96 well plate in triplicate and repeated at least twice. OD_600_was measured at specific time points for 18-24 h and the following analyses were performed 1) mutant OD_600_values were compared to the WT JE2 strain after overnight growth and 2) the growth ratio at each time point was calculated using OD_600_(mutant) ÷ OD_600_(WT). Mutants with a significantly lower OD_600_compared to WT JE2 after overnight growth or a growth ratio <0.5 at both 2 h as well as 4 h points were selected, and transposon insertion mutants were used to transduce the JE2 and MRSA 1369 background strains using lysates made from phage 11, as described above. Growth assays for these newly generated mutants in both JE2 and MRSA 1369 backgrounds were performed in BHI and HU for 24h. Based on the results of these assays, four mutants were selected targeting TCA cycle (Δ*fumC*, Δ*sucD*), pyruvate oxidation (Δ*lpdA*), and mannitol metabolism (Δ*mtlD*) in the uropathogenic MRSA 1369 strain background for further examination.

### Growth Kinetics of WT and mutants

Overnight cultures of MRSA 1369 and mutant strains were pelleted and washed in D-PBS. Bacteria were then inoculated 1:1000 in pooled HU or BHI and incubated at 37°C without shaking for 24 h. At 0, 2, 4, 6, 8, and 24 h time points, a 10 μl sample was dilution plated to determine CFU/ml on tryptic soy broth (TSA) plain or TSA containing 10 μg/mL of erythromycin for transposon mutants where appropriate.

For each strain, the doubling time (DT) was calculated:

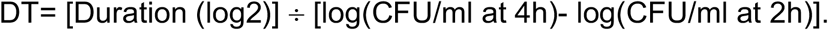

### In vitro competition in HU

Mid-log phase cultures of MRSA 1369 WT and mutant strains were washed in D-PBS and inoculated in pooled HU at 1:1 ratio (2×10^6^ CFU/ml each for the WT and the mutant). The WT and mutant CFUs at 0 h were differentially enumerated by dilution plating the inoculum on plain TSA and TSA containing 10 μg/ml erythromycin for total and mutant CFU/ml, respectively. After incubation at 37°C, samples were taken at 4 and 24 h to determine the number of WT and mutant CFUs recovered by dilution plating on TSA and TSA containing 10 μg/ml erythromycin. For 0 (inoculum), 4, and 24 h, WT CFU/ml = Total CFU/ml – mutant CFU/ml. The competitive index (CI) for each mutant strain was calculated at 4 h and 24 h time points:

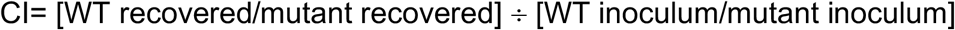

### Mouse model of CAUTI

As approved by the Institutional Animal Care and Use Committee (IACUC) at the University of Louisiana at Lafayette (protocol number 2020-8717-025), we experimentally induced CAUTI in 8-to 10-week-old female C57BL/6 mice, as described previously (12, 22). In brief, the overnight cultures of MRSA 1369 WT and select mutant strains were inoculated 1:10 in fresh BHI medium and incubated at 37°C shaking at 200 rpm. Mid-log phase (OD_600_= 0.6) cultures were then pelleted, washed, and resuspended in D-PBS. Inoculum of 50μl (equivalent to ∼5×10^7^CFU/ml) was inoculated immediately after implantation of a 4-to 5-mm piece of silicone tubing (catheter) via transurethral insertion into the bladder of anesthetized mice. The catheter implant remains in the urine bladder during infection.

For single infections, MRSA 1369 WT and mutant strains were inoculated separately into the urinary bladders of anesthetized mice. Mice were sacrificed at 24 hpi and spleen, kidneys, bladder, and catheter implant were aseptically and sequentially harvested. The CFUs were determined by dilution plating the homogenates of bladder, combined halves of bisected left and right kidneys, and spleen. The catheter implant recovered from the bladder was vortexed in 1ml sterile D-PBS for 1 minute to recover bacteria, which were then enumerated by dilution plating. Statistical significance between the organ burdens from WT- and mutant-infected mice were determined using Mann Whitney U test.

For *in vivo* competition experiments, mid-log phase cultures of MRSA WT and mutant were mixed 1:1 to obtain 5 × 10^7^ CFU/ml total inoculum resuspended in 50μl sterile D-PBS. Anesthetized female C57BL/6 mice were then inoculated via transurethral catheterization, as described above. The inoculum was dilution plated on plain TSA plates to enumerate total inoculum (WT + mutant CFU/ml) and on TSA + 10 μg/ml erythromycin to enumerate mutant inoculum CFU/ml. After overnight incubation at 37°C, colonies were counted. WT inoculum = total inoculum – mutant inoculum. At 24 hpi, organs and catheter from infected mice were harvested and processed as described above. These were dilution plated either on plain TSA (total CFU/ml recovered) or on TSA+ erythromycin (mutant CFU/ml recovered). WT recovered = Total recovered – mutant recovered. Competitive index (CI) was calculated:

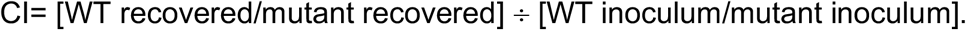

CI= 1 represents that both WT and mutant strains are equally competitive in the mouse UT; CI<1 represents that the mutant has competitive advantage over WT for colonizing the mouse UT; CI>1 represents that WT has competitive advantage over the mutant.

### Hydrogen peroxide (H_2_O_2_) killing assay

Overnight cultures of MRSA 1369 WT and mutant strains in BHI were centrifuged, washed in D-PBS, and resuspended at 1:100 dilutions in either fresh BHI medium or HU supplemented with 15mM H_2_O_2_. Inoculum CFUs were determined by dilution plating. Cultures were incubated at 37ºC and 200rpm shaking and bacterial CFUs were determined by dilution plating at 4 h.

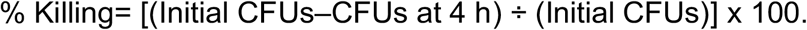

### Cell surface hydrophobicity assay

Cell surface hydrophobicity was determined using MATH (microbial adhesion to hydrocarbon assay) (16) where mid log-phase culture of MRSA 1369 WT and mutant strains (OD_600_=0.6 in BHI) were centrifuged, washed in D-PBS, and exposed to HU for 2 h. MRSA strains in HU were centrifuged, washed, and resuspended in sterile D-PBS to OD_600_of 0.5 and plated for inoculum CFU (C_i_) enumeration. One ml of the bacterial suspension was mixed with 125μl hexadecane, vortexed for 1 min and incubated at room temperature for 30 min. The CFUs in the aqueous phase (C_aq_) were then enumerated by dilution plating. %Hydrophobicity = [(C_i_– C_aq_)÷(C_i_)] x 100

### Whole blood killing assay

The mid log-phase cultures of MRSA 1369 WT and mutant strains (OD_600_=0.6 in BHI) were centrifuged, washed in D-PBS, and exposed to HU for 2 h. Next, 200 μl of hirudin-anticoagulated human blood was mixed with 2×10^6^ CFUs of the mutant or WT MRSA strains in a 96-well plate. After incubation at 37°C and continuous shaking, samples were extracted at 1 h or 4 h to enumerate surviving bacteria by dilution plating for CFUs.

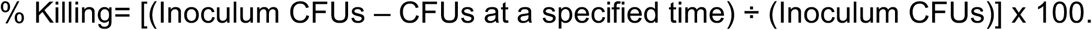

### Statistical analysis

Statistical tests were performed using Prism 9.0 (www.graphpad.com). Data from multiple biological replicates with two or more technical replicates and two or more biological replicates for each experiment were pooled together. Error bars in the figures represent standard error of the mean. Competitive indices for each mutant strain compared to WT in vitro or in vivo were analyzed using one sample t- and Wilcoxon test against a theoretical median of 1. Growth, doubling time, hydrophobicity, and organ burden data from single infection experiments were compared using Mann-Whitney U statistic. Dissemination rates were compared using Bernard’s Test. Data were considered statistically significant if P ≤ 0.05.

## Results

### Genome-wide screen for MRSA genes required for growth in HU

We screened MRSA JE2 and the corresponding 1920 transposon mutants from the Nebraska transposon mutant library (NTML) for growth in HU by measuring OD_600_over an overnight time course. The growth ratios= OD_600_(mutant)/OD_600_(WT) calculated at each time point are presented as heat map in Fig 1A; raw OD_600_values are provided as supplementary file S1. Of these, sixteen mutants in the JE2 background showed significant growth defects in HU (defined as growth ratio≤0.5 at 2 and 4 h) and were matched with MRSA 1369 gene expression data following 2 h exposure to HU (Fig 1B and previously published (16)). Of these sixteen mutants, *fumC, sucD*, and *mtlD* expression was significantly upregulated in MRSA 1369 following 2 h exposure to HU. Thus, Δ*fumC*, Δ*sucD* (defective in TCA cycle enzymes), Δ*lpdA* (defective in pyruvate oxidation enzyme), and Δ*mtlD* (defective in mannitol metabolism) mutants were selected for further analysis. Additionally, while *lpdA* expression was not altered in MRSA 1369 following 2 h exposure to HU, the mutant displayed the most severe growth defect in HU. Thus, Δ*lpdA* (defective for dihydrolipoamide dehydrogenase activity) was also selected for further analysis. Next, we used phage transduction to move the corresponding transposon mutants into the MRSA 1369 strain, which is a clinical strain isolated from HU.

**Figure 1.**
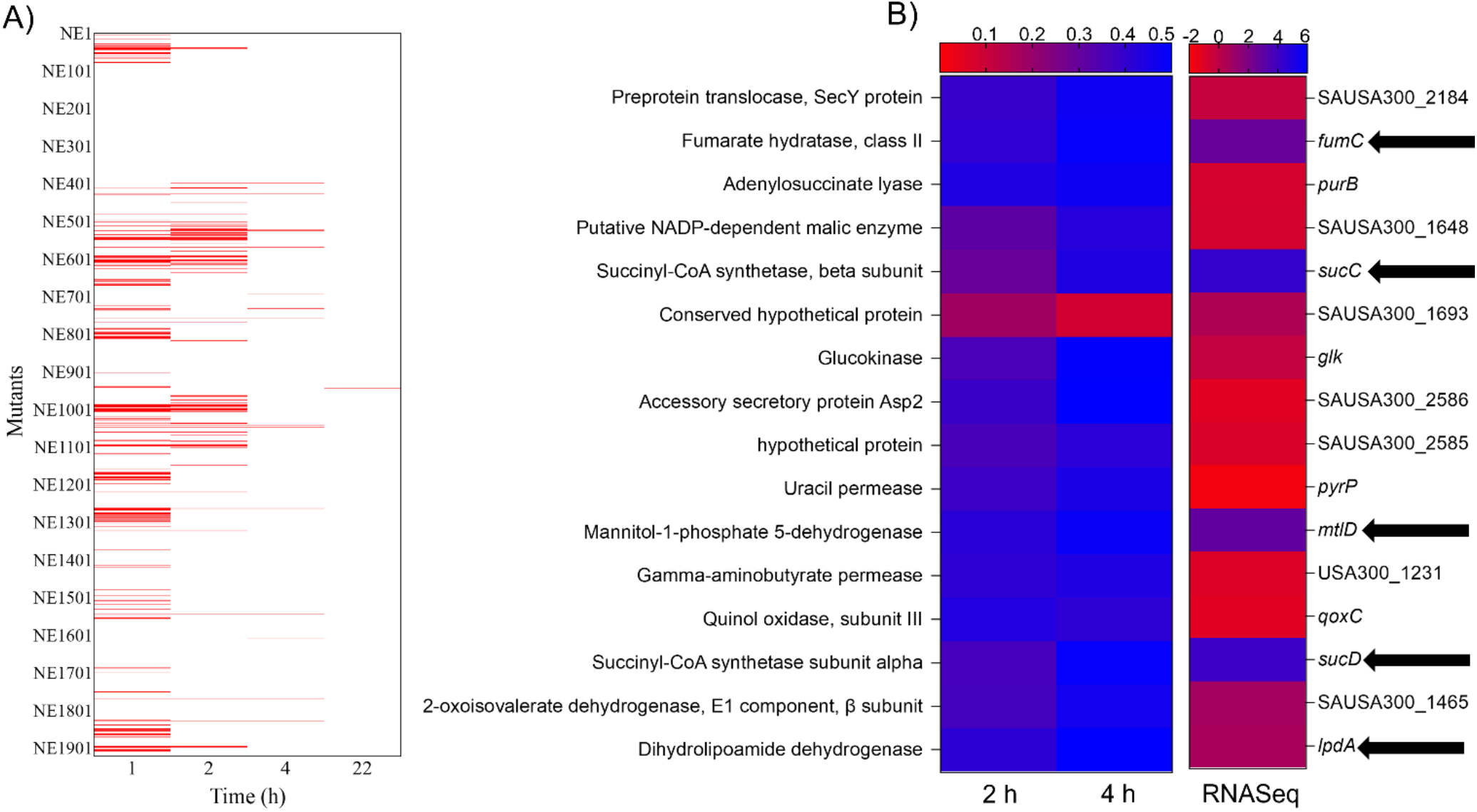
Screening of MRSA JE2 mutants for growth and MRSA 1369 gene expression in human urine (HU). MRSA JE2 WT and 1920 mutant strains from the Nebraska transposon mutant library (NTML) were screened for the overnight growth in HU by measuring OD_600_. A) The heat map shows growth ratio (OD_600_mutant/OD_600_WT) for all strains in the library. All mutants with a ≤0.5 ratio are denoted in red. B) Of strains with significant growth defects in HU (growth ratio≤ 0.5 at 2 and 4 h), these were matched with gene expression data from MRSA 1369 WT strain exposed to HU for 2 h. The genes with observed correlation between defective growth ratio and upregulation of gene expression are indicated by arrows. The growth curves were repeated at least three times, each with three technical replicates. Gene expression results are from our previously published RNASeq analysis (16).

### WT and mutant MRSA 1369 *in vitro* growth kinetics in HU

To validate that the identified JE2 mutants are also defective for growth in HU in the MRSA 1369 background, but displayed normal growth in a nutrient rich medium, growth curves were repeated in BHI and HU (Fig 2). All MRSA 1369 mutant strains displayed similar growth in BHI compared to WT MRSA 1369 (Fig 2A). In HU, only Δ*lpdA* showed 4-fold lower CFUs (2.8×10^7^CFU/ml) compared to that of WT (1.2×10^8^CFU/ml) at 24 h. All other mutants, including Δ*sucD* (9.1×10^7^CFU/ml), *fumC* (9.6×10^7^CFU/ml), and Δ*mtlD* (5.3×10^7^CFU/ml), displayed similar CFUs at 24 h compared to WT (Fig 2B). Additionally, all isogenic mutant strains displayed similar growth kinetics in BHI compared to WT MRSA 1369, with doubling times around 20 min (Fig 2C). Lastly, the growth kinetics of MRSA 1369 Δ*sucD*, Δ*fumC*, and Δ*mtlD* were not significantly different from that of WT as emphasized by the similar doubling times of Δ*sucD* (52±15 min), *fumC* (46±17 min), Δ*mtlD* (61±25 min), and WT (46±12 min) (Fig 2D). These results suggest that the TCA cycle and mannitol metabolism are not required for growth in HU. In contrast, MRSA 1369Δ*lpdA* displayed a significant defect for growth in HU with a doubling time (224±37 minutes) that was 5-fold higher than that of WT (Fig 2D). Thus, acetyl CoA synthesis appears to play an essential role in the growth and survival of MRSA 1369 in HU.

**Figure 2.**
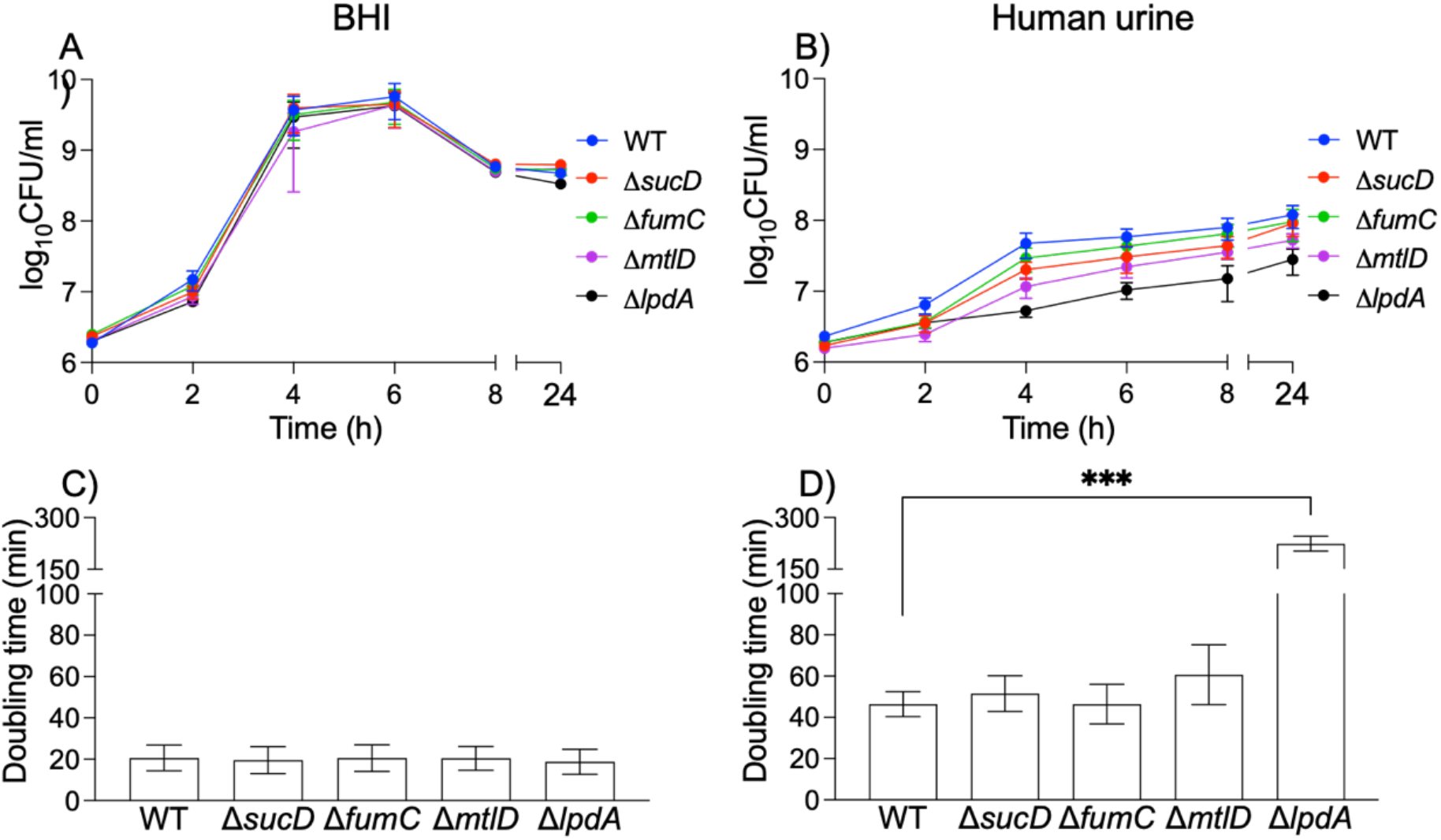
Growth curves of MRSA 1369 WT and select mutants. MRSA strains were grown at 37°C, static, for 24 h in either nutrient-rich BHI (A and C) or pooled human urine (B and D) and CFU/ml were determined by dilution plating on TSA at specific time points. The growth of WT and mutant MRSA strains are presented as growth curves showing average CFU/ml ± standard error of mean (A and B) and as average doubling time ± standard deviation (C and D). The data are from 2 biological replicates for BHI and from 3 to 4 biological replicates for human urine; each biological replicate had 2 technical replicates. The doubling time for each mutant was compared with the WT using unpaired t test. *** *P*≤0.001 compared to WT control.

### *In vitro* competition between WT and mutant MRSA 1369 in HU

Next, we set up in vitro co-culture experiments by inoculating equal numbers of a MRSA 1369 WT and mutants in HU. The direct competition for amino acids and peptides in the HU makes the urinary fitness advantage that WT or mutants may have over one another more discernible. We enumerated WT and mutant CFUs and calculated separate competitive index (CI) values at 4 h (Fig 3A) and 24 h (Fig 3B) of co-culture. Similar to the results from the growth curve assays, Δ*fumC* was just as fit as WT for growth in HU at either 4 or 24 h. Additionally, Δ*lpdA* displayed a significant competitive disadvantage compared to WT for growth in HU at both 4 and 24 h time points. In contrast, to the growth curve experiments, Δ*sucD* appeared to be at competitive disadvantage at 4 h, although by 24 h it appeared to be competitively equal with WT for the growth in HU. Additionally, Δ*mtlD* displayed a significant competitive disadvantage at both 4 and 24 h time points.

**Figure 3.**
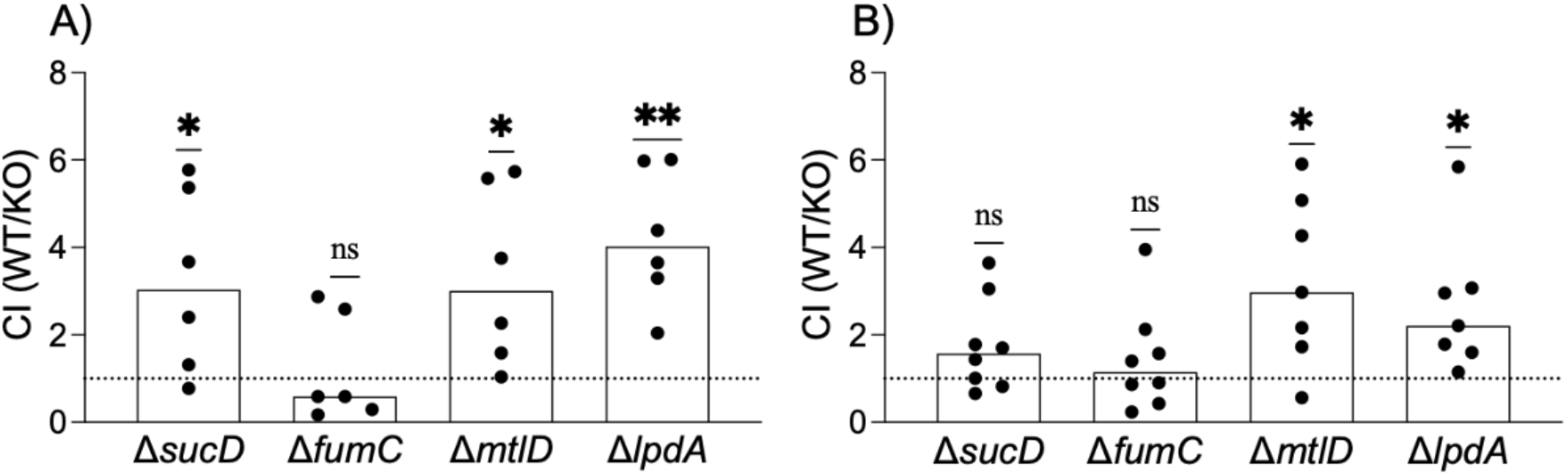
Competition between WT and mutant MRSA 1369 strains for *in vitro* growth in pooled human urine (HU). WT MRSA 1369 was separately co-cultivated with each gene mutant at 1:1 ratio in pooled HU at 37°C, static. Competitive index (CI) values for each mutant strain (KO) were calculated by enumerating WT and KO CFUs recovered at (A) 4h and (B) 24h post-inoculation. CI values are shown as scatter plots with each point representing a technical replicate and median as the measure of central tendency. CI=1, shown as a dotted line, represents that WT and KO are equally competitive, CI<1 denotes that the KO has a competitive advantage over WT, and CI>1 indicates that the WT has a competitive advantage over the KO. Data from 3 or 4 biological replicates, each with 2 technical replicates are shown. Data were compared against a theoretical median of 1 using one sample t test and Wilcoxon test. **, *P*≤0.01; *, *P*≤0.05.

### WT and mutant MRSA 1369 *in vivo* fitness in the mouse model of CAUTI

To assess the role of metabolism and amino acid synthesis during infection, we used *in vivo* competition experiments in the murine model of CAUTI. We co-infected 6-8 weeks old, female C57BL/6 mice with equal numbers of MRSA 1369 WT and individual mutant strains (1:1 inoculum of WT and mutant). At 24 hpi, the bacterial burden in the bladder, kidneys, and catheter were determined by dilution plating and competitive index (CI) values were calculated (Fig 4). Both the TCA cycle enzyme mutants Δ*sucD* (Fig 5A) and Δ*fumC* (Fig 5B) were less fit for UT colonization as indicated by significantly higher CI values for the bladder (median CI for Δ*sucD*= 5.6, Δ*fumC*= 1.8) and the kidneys (median CI for Δ*sucD*= 10, Δ*fumC*= 3.4). Additionally, while Δ*sucD* was less fit in colonizing the catheter implant compared to WT (median CI= 5.3), Δ*fumC* (median CI= 0.8) was not (Fig 4A, B). Additionally, kidney infection was detected in 8/11 WT and Δ*sucD* co-infected mice and 7/11 WT and Δ*fumC* co-infected mice. The Δ*mtlD* mutant was also less fit for colonizing the UT (median CI for bladder= 3.8; catheter implant= 3.3; kidneys= 3.5); kidney infection was also detected in 6/8 mice co-infected with WT and Δ*mtlD* (Fig 4C). Consistent with the in vitro data, Δ*lpdA* was less fit in colonizing the UT as indicated by significantly higher CI values compared to WT for the bladder (median CI= 15.5) and the catheter implant (median CI= 11.4). Notably, only 2/8 WT and Δ*lpdA* co-infected mice had detectable kidney infection (Fig 4E). Overall, these results indicate that Δ*sucD* and Δ*lpdA* mutants display the largest fitness defects in UT colonization in competition experiments.

**Figure 4.**
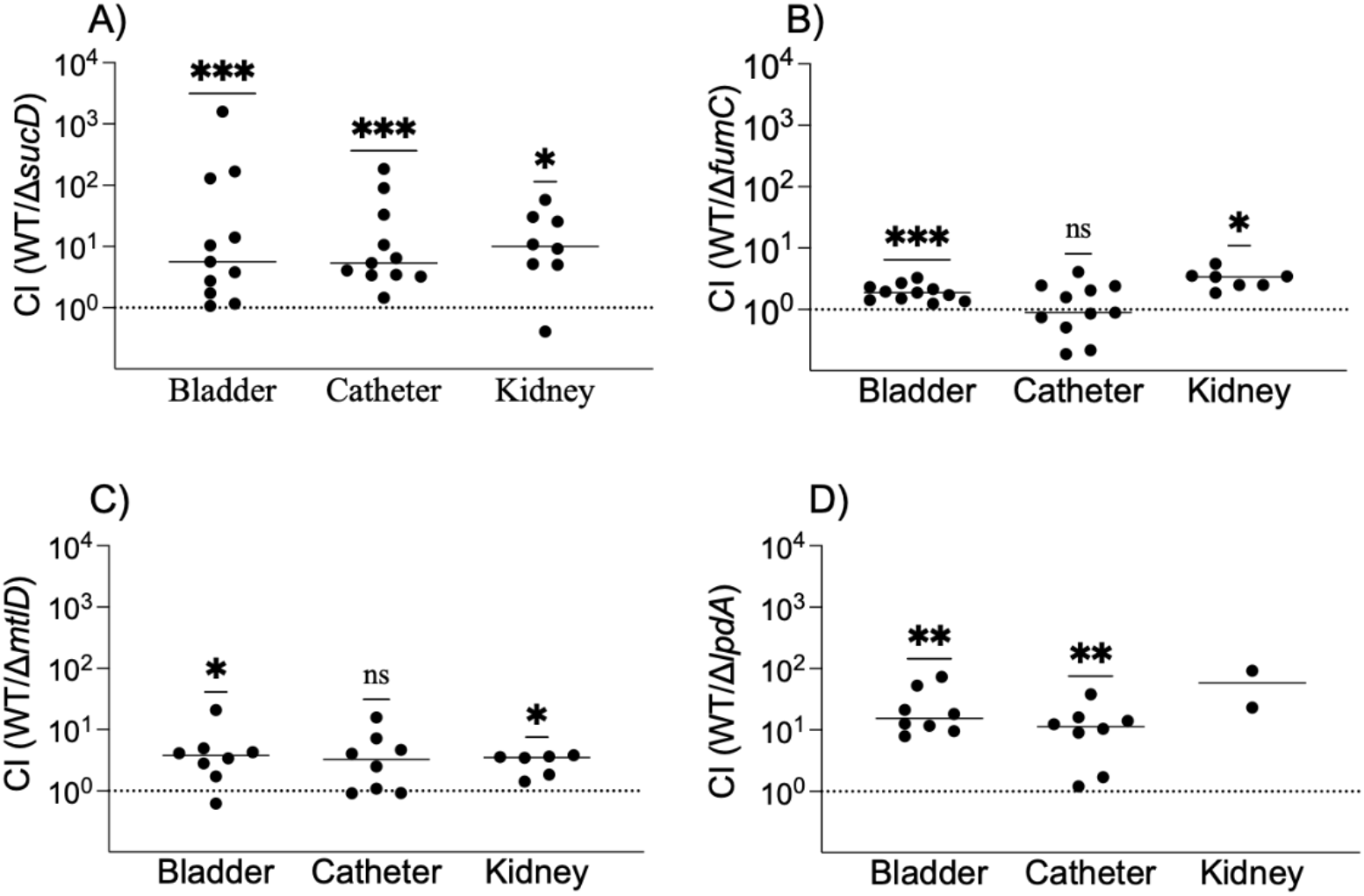
*In vivo* competition examining the fitness of MRSA 1369 mutants compared to the WT. C57BL/6 female mice were catheterized and then inoculated transurethrally with 1:1 mixture of MRSA 1369 WT and either A) Δ*sucD*, B) Δ*fumC*, C) Δ*mtlD*, or D) Δ*lpdA* mutants. At 24 hpi, WT and mutant CFU burden in bladder, kidneys, and catheter was determined. Competitive index (CI) values for the urinary bladder, kidneys, or catheter from each mouse are presented as scatter diagrams with median as the measure of central tendency. CI=1, shown as a dotted line, represents that WT and mutant are equally competitive, CI<1 denoted that the mutant has a competitive advantage over WT, and CI>1 indicates that the WT has a competitive advantage over mutant. Data from ≥2 biological replicates were compared against a theoretical median of 1 using one sample t test and Wilcoxon test. ***, P≤0.001; **, P≤0.01; *, P≤0.05; ns, not significant.

**Figure 5.**
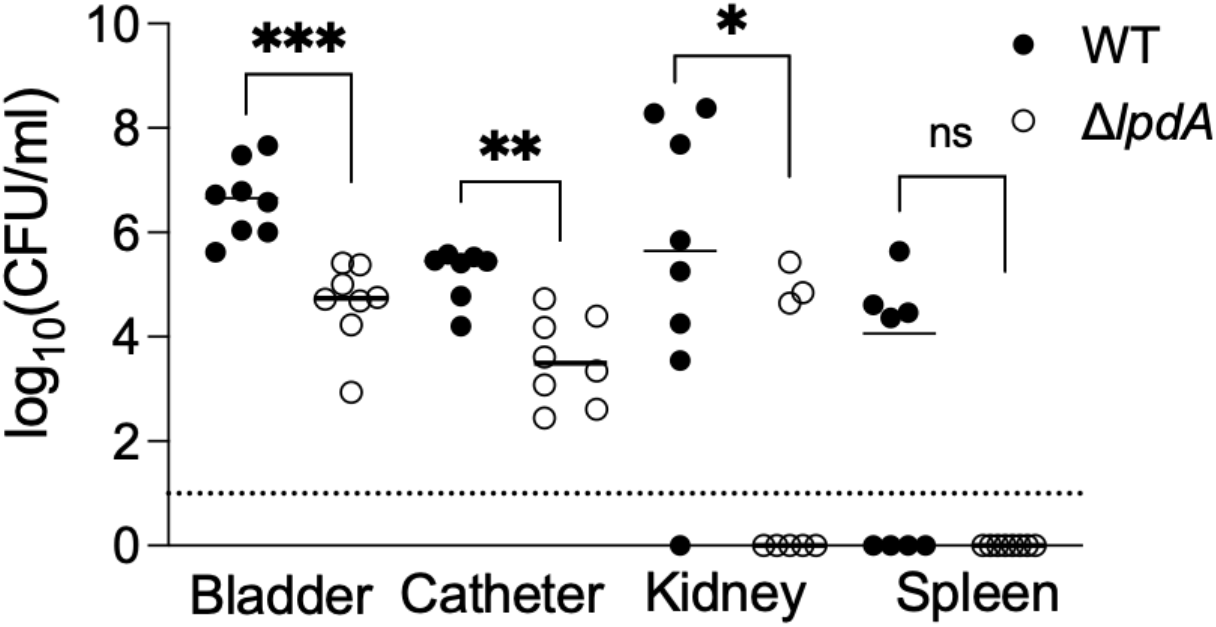
Urinary pathogenesis of Δ*lpdA* in an *in vivo* mouse model of CAUTI. C57BL/6 female mice were inoculated with WT or Δ*lpdA*. At 24 hpi, WT and Δ*lpdA* CFUs in the spleen, bladder, kidneys, and catheter were determined. The data are presented as scatter diagrams showing organ burden from an individual mouse and median as the central tendency. Dotted line represents limit of detection (LOD) of 10CFU/ml. Data from 8 mice per group from ≥2 biological replicates were compared using Mann-Whitney U test; ***, P≤0.001; **, P≤0.01; *, P≤0.05; ns, not significant.

### Uropathogenesis of Δ*lpdA* during mono-culture CAUTI

The results from the *in vivo* competition experiments prompted us to further examine the uropathogenesis of Δ*lpdA* via mono-culture infection using the mouse model of CAUTI. The Δ*lpdA* mutant displayed significantly lower CFUs in the bladder (Δ*lpdA* median= 55,000 CFU/ml, WT= 45.5×10^5^ CFU/ml; *P*=0.0002), the catheter implant (Δ*lpdA* median= 3150 CFU/ml, WT= 2.8×10^5^ CFU/ml; *P*=0.0012), and the kidney (Δ*lpdA* < LOD, WT= 4.4×10^5^ CFU/ml; *P*=0.037) CFUs compared to WT infected mice (Fig 5). Notably, the dissemination rate to the spleen was significantly higher among mice infected with WT (4/8) compared to those infected with Δ*lpdA* (0/8). Overall, these results revealed that Δ*lpdA* plays a role in the colonization of the lower UT as well as in dissemination to the kidneys and the spleen in a mouse model of CAUTI.

### Survival of Δ*mtlD* and Δ*lpdA* mutants in the presence of H_2_O_2_

Activity of MtlD and LpdA enzymes regulate *S. aureus* reactive oxygen species (ROS) stress response either by replenishing intracellular mannitol reserves (23) or by regulating membrane fluidity (17, 19). To investigate the potential mechanisms by which these genes promote infection, we examined Δ*mtlD* and Δ*lpdA* for survival in the presence of H_2_O_2_, a bactericidal product of activated macrophages and neutrophils. In nutrient rich BHI, both Δ*mtlD* and Δ*lpdA* mutants were significantly more susceptible to H_2_O_2_ at 4 h (Fig 6A). Notably, in HU only Δ*mtlD* was significantly more sensitive to H_2_O_2_ at 4 h (Fig 6B).

**Figure 6.**
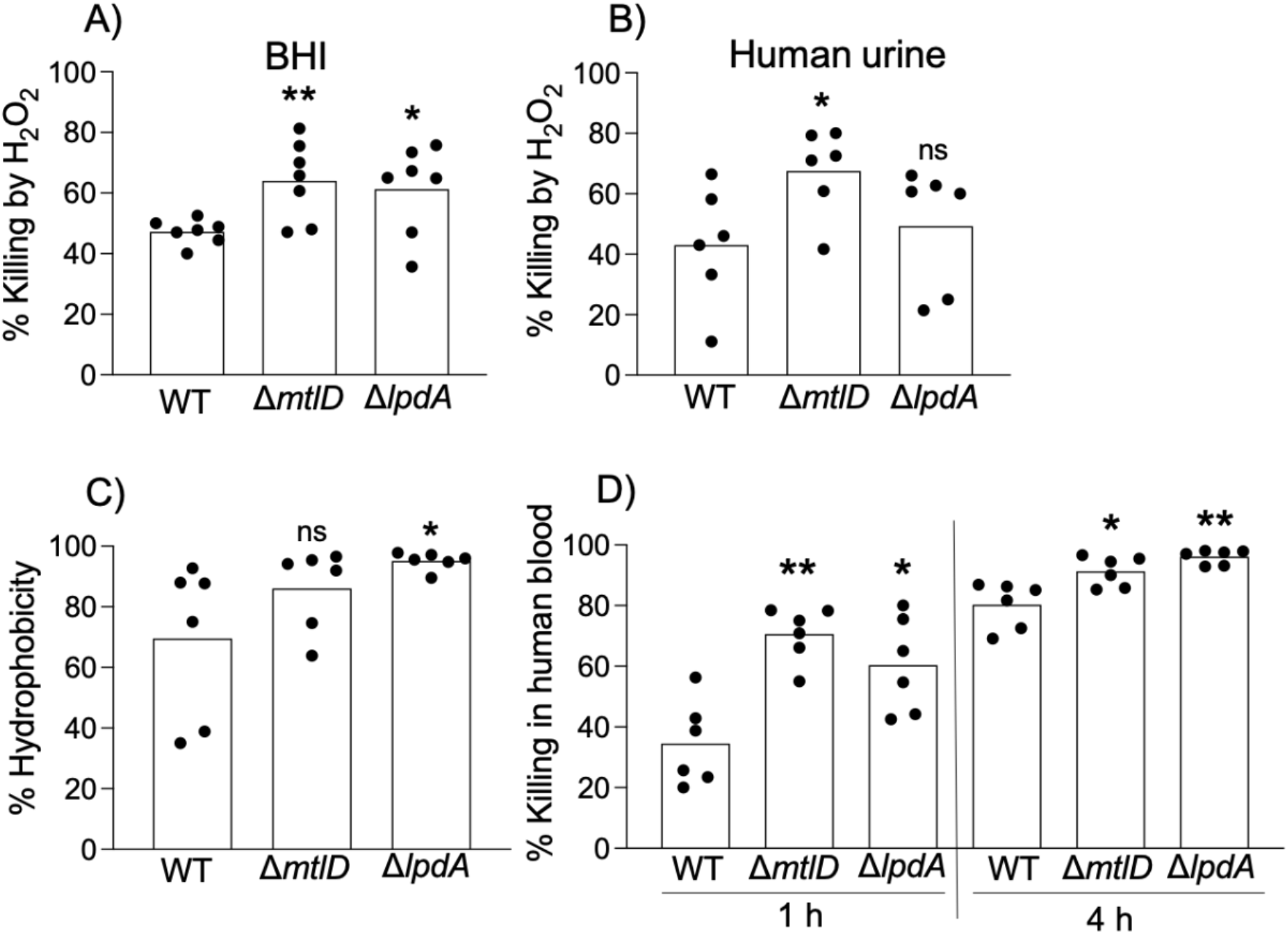
Δ*mtlD* and Δ*lpdA* susceptibility to H_2_O_2_, cell surface hydrophobicity, and in vitro killing by whole blood. WT, Δ*mtlD*, and Δ*lpdA* were exposed to 15 mM H_2_O_2_ for 4 h in either A) BHI or B) pooled human urine (HU). The inoculum (0 h) and surviving CFUs after 4 h-long exposure to H_2_O_2_ were determined. Additionally, following 2 h exposure to urine, WT and Δ*lpdA* were compared for (C) cell surface hydrophobicity, (D) % killing in human blood. Scatter plots show individual technical replicates from 3 biological replicates. Histograms represent average for biological replicates. The data for each mutant were compared with the WT (at 1 h or 4 h time point for % killing in human blood) using unpaired T test; **, P≤0.01; *, P≤0.05; ns, not significant.

### Δ*lpdA* and Δ*mtlD* cell surface hydrophobicity

The essential role Δ*lpdA* plays in vivo led us to further assess the how LpdA contributes to virulence. First, we assessed membrane fluidity, as LpdA can modulate membrane fluidity and surface hydrophobicity of *S. aureus* through the regulation of branched chain fatty acid synthesis (33–35). Using MATH (microbial adhesion to hydrocarbon assay) protocol, we compared WT and Δ*lpdA* strains grown in HU for 2 h for their ability to adhere to the hydrocarbon, hexadecane. We observed a significant increase in the hydrophobicity of Δ*lpdA* compared to the WT (Fig 6C). The hydrophobicity of Δ*mtlD* was not significantly affected (Fig 6C), which suggests that the increase in hydrophobicity is directly caused by a lack of LpdA.

### Survival of Δ*lpdA* and *ΔmtlD* in human blood

MRSA 1369 typically disseminates to the blood and spleen in ∼50% of infected mice at 24 hpi during CAUTI (12). Consistently, in this study, 4/8 (50%) WT-infected mice displayed high CFUs in the spleen (Fig 5). In contrast, the absence of infection in the spleen in any of the mice singly infected with *ΔlpdA* at 24 hpi (Fig 5) suggests that *ΔlpdA* may be defective for survival in blood. Indeed, compared to the WT, *ΔlpdA* was significantly more susceptible to killing in whole human blood at both 1 h and 4 h following growth in urine (Fig 6D). Furthermore, *ΔmtlD* was also more susceptible to killing in whole human blood at both 1 h and 4 h following growth in urine (Fig 6D). This data suggests *ΔlpdA* and *ΔmtlD* provide resistance to stress encountered in the UT and during dissemination during CAUTI.

## DISCUSSION

First detected in hospitalized patients in the 1960s, MRSA has spread in the community for the last three decades as a commensal colonizing the human skin, nasopharynx, the lower digestive tract (24). MRSA colonization significantly increases the risk of infections ranging from moderately severe skin infections to potentially fatal pneumonia and sepsis (25). In addition, MRSA has also emerged as an important etiology of hospital-acquired UTIs, primarily arising from the contamination of indwelling urinary catheters (26). Studies using the mouse model of ascending UTI have established that MRSA can colonize murine UTs either in the presence (complicated UTI) or the absence (uncomplicated UTI) of a catheter and specific virulence traits central to MRSA uropathogenesis have been identified (12, 27). Notably, the contribution of MRSA metabolism to its survival and fitness in the UT has not been deciphered, despite the known role of amino acid utilization via TCA cycle and gluconeogenesis in the *in vivo* urinary fitness of both UPEC and *P. mirabilis* (13-15). Specifically, FumC, the fumarase enzyme, of the TCA cycle has been reported to be required for the colonization of the UT by UPEC and *P. mirabilis* (15). Furthermore, FumC contributes to respiratory survival and persistence of *S. aureus* (28). Interestingly, UT colonization by *P. mirabilis* also requires the activity of glycolysis and Entner-Dodoroff pathways catabolizing glucose into pyruvate, while these glucose utilization pathways are dispensable for the *in vivo* fitness of UPEC (14). The essentiality of the TCA cycle and gluconeogenesis is unsurprising as the UT in non-diabetic humans is a nutrient-poor, carbohydrate-lacking microenvironment where short peptides and amino acids in urine are the principal carbon sources available to the colonizing uropathogens. We recently reported that exposing MRSA 1369 to HU induces the expression of genes encoding oligopeptide import systems and TCA cycle enzyme, while suppressing the expression of glycolysis genes (16). Overall, these observations indicate that different uropathogens rely on divergent central carbon metabolism pathways for successful colonization of the UT.

To further investigate the role of metabolism in MRSA uropathogenesis, we screened a MRSA transposon insertion mutant library and identified 16 mutants that were defective for growth in HU but grew normally in nutrient rich BHI. Of these, we selected mutants ablated for enzymes catalyzing TCA cycle, succinyl-coA synthase (Δ*sucD*) and fumarase (Δ*fumC*); mannitol metabolism, mannitol-1-phosphate dehydrogenase (Δ*mtlD*); and pyruvate oxidation, dihydrolipoamide dehydrogenase (Δ*lpdA*), for *in vitro* and *in vivo* competition assays. SucD catalyzes substrate level phosphorylation via the hydrolysis of succinyl CoA into succinate and GTP (29). In *E. coli*, the FumC is an iron-independent fumarase enzyme that catalyzes conversion of fumarate to malate in the oxidative TCA cycle (30, 31). MtlD catalyzes the reversible conversion of fructose-6-phosphate to mannitol-1-phosphate, a key step in mannitol metabolism. Lastly, LpdA catalyzes pyruvate oxidation to acetyl-coA that bridges glycolysis and the TCA cycle. Thus, many of the mutations that displayed growth phenotypes or competitive fitness defects following growth in HU, were essential enzymes in metabolism, and specifically in the TCA cycle.

For successful UT colonization, MRSA must derive carbon by catabolizing amino acids and short peptides in urine; the carbon is then used to fuel the TCA cycle. We have previously reported the upregulation of TCA cycle genes including *sucD* and *fumC* in the MRSA1369 strain exposed to HU for 2 h (16). Here, we show that compared to WT MRSA 1369, neither Δ*sucD* nor Δ*fumC* is significantly defective for growth in HU, which differed from the JE2 background mutants, nor did they display a competitive disadvantage over WT. The biochemical pathway used by the Δ*sucD* and Δ*fumC* mutants for growth in HU is unclear as *S. aureus* lacks genes encoding isocitrate lyase and malate synthase, which can bypass the TCA cycle via glyoxalate in the absence of *sucD* and *fumC* (32). Interestingly, however, when competed with the WT MRSA 1369 strain in an *in vivo* mouse model of CAUTI, Δ*sucD* and Δ*fumC* were found to be required for fitness (median CI< 10 compared to WT), thus confirming the requirement of a functional TCA cycle for MRSA in urinary pathogenesis.

The enzyme mannitol-1-phosphate dehydrogenase *aka* M1PDH (encoded by *mtlD*) catalyzes the reversible conversion of fructose-6-phosphate to mannitol-1-phosphate, a key step in mannitol metabolism. Dephosphorylation of mannitol-1-phosphate can replenish the intracellular reserves of mannitol, which alleviates *in vivo* stress due to high ROS concentrations and acidic pH and facilitates MRSA fitness in mouse models of systemic infection (23). We observed that Δ*mtlD* was at a significant competitive disadvantage over WT for in vitro growth in HU as well as for in vivo UT colonization (median CI< 10 compared to WT). These phenotypes may be attributed to the increased sensitivity of Δ*mtlD* mutant to H_2_O_2_ and whole blood *in vitro*. While humans do not produce mannitol and the baseline urine mannitol concentration is very low (33), consumption of mannitol sweeteners can raise urine mannitol levels to 14 mg/ml. Urine mannitol can be fermented by MRSA MtlD to fructose-6-phophate which can be further processed through the energy metabolism pathway (glycolysis) although the physiological relevance of this pathway during UTI would depend on the dietary exposure to mannitol sweeteners. These data indicate that mannitol metabolism plays an important role in MRSA UT fitness via increased sensitivity to H_2_O_2_ and blood.

Dihydrolipoamide hydrogenase encoded by *lpdA* is one of the four constituent enzymes of branched-chain keto acid dehydrogenase (BKD) multi-enzyme complex. BKD enzymes catalyze pyruvate oxidation to acetyl-coA that bridges glycolysis and the TCA cycle. Acetyl-coA is also a precursor for the synthesis of branched- and straight-chain fatty acids (BCFA and SCFA), the principal determinants of staphylococcal cell membrane fluidity (34). An *lpdA* (BKD deficient) mutant strain of *S. aureus* has been observed to exhibit decreased membrane fluidity, increased susceptibility to H_2_O_2_, and reduced survival in a murine model of systemic infection (17). Our previous report indicated a 2 h-long exposure to HU did not alter MRSA *lpdA* transcription, which suggests that at least at early time points pyruvate oxidation is not needed for MRSA growth in a glucose-poor growth medium such as healthy HU (16). We report here that Δ*lpdA i)* grew significantly slower than the WT MRSA 1369 and JE2 strains in urine from healthy volunteers; *ii)* was severely defective compared with the WT in an *in vivo* competition mouse model of CAUTI (median CI>10); *iii)* was defective in a mouse model of CAUTI; and *iv)* displayed significantly increased cell surface hydrophobicity and susceptibility to killing in blood following a 2 h-long *in vitro* exposure to HU. Together, these data suggest that the cell surface of Δ*lpdA* is altered, which may in turn affect the fitness of MRSA in the UT, as it results in increased hydrophobicity and sensitivity to killing in whole blood. Additionally, the increased sensitivity to whole blood killing may explain why the Δ*lpdA* mutant was not recovered from the splenic homogenates of the mouse model of CAUTI at 24 hpi.

Overall, our results dictate the importance of the TCA cycle, mannitol metabolism, and synthesis of branched chain fatty acid for the fitness of MRSA in the UT environment. The continual emergence of antibiotic-resistant strains resulting from the acquisition of mobile genetic elements remain the major challenge in the successful treatment of MRSA infections. A better understanding of the metabolic pathways utilized in the host UT for the survival and proliferation may help us develop novel therapeutics against MRSA UTI.

## Acknowledgements

We would like to thank BEI Resources for providing the Nebraska Transposon Mutant Library (NR-48501). This work was supported by grants R21 AI165939 (to RK) from the NIH National Institute of Allergy and Infectious Diseases and K01-DK128381-01A1 (to JNW) and R01-DK051406 (to CLPO and SJH) from the NIH National Institute of Diabetes and Digestive and Kidney Diseases.

## Conflicts of Interests

The authors declare no conflicts of interest

